# Inferring the ancestry of parents and grandparents from genetic data

**DOI:** 10.1101/308494

**Authors:** Jingwen Pei, Yiming Zhang, Rasmus Nielsen, Yufeng Wu

**Affiliations:** Department of Computer Science and Engineering, University of Connecticut, Storrs, CT, U.S.A; Departments of Integrative Biology and Statistics, University of California, Berkeley, Berkeley, CA, U.S.A; Museum of Natural History, University of Copenhagen, Copenhagen K, 1350, Denmark

## Abstract

Inference of admixture proportions is a classical statistical problem in population genetics. Standard methods implicitly assume that both parents of an individual have the same admixture fraction. However, this is rarely the case in real data. In this paper we show that the distribution of admixture tract lengths in a genome contains information about the admixture proportions of the ancestors of an individual. We develop a Hidden Markov Model (HMM) framework for estimating the admixture proportions of the immediate ancestors of an individual, i.e. a type of decomposition of an individual’s admixture proportions into further subsets of ancestral proportions in the ancestors. Based on a genealogical model for admixture tracts, we develop an efficient algorithm for computing the sampling probability of the genome from a single individual, as a function of the admixture proportions of the ancestors of this individual. This allows us to perform probabilistic inference of admixture proportions of ancestors only using the genome of an extant individual. We perform extensive simulations to quantify the error in the estimation of ancestral admixture proportions under various conditions. To illustrate the utility of the method, we apply it to real genetic data.

**Author summary:** Ancestry inference is an important problem in genetics and is used commercially by a number of companies affecting millions of consumers of genetic ancestry tests. In this paper, we show that it is possible, not only to estimate the ancestry fractions of an individual, but also, with some uncertainty, to estimate the ancestry fractions of an individual’s ancestors. For example, if an individual traces his/her ancestry 50% to Asia and 50% to Europe, it is possible to distinguish between the individual having two parents that each are 50:50 composites of Asian and European ancestry, or one parent from Asia and one from Europe. It is likewise also possible to make inferences about grandparents. We present a computationally efficient method for making such inferences called PedMix. PedMix is based on a probabilistic model for the descendant and the recent ancestors. PedMix infers admixture proportions of recent ancestors (parents, grandparents or even great grandparents) using whole-genome genetic variation data from a focal individual. Results on both simulated and real data show that PedMix performs reasonably well in most scenarios.

## Introduction

Ancestry inference is one of the most commonly used tools in human genetics. It arguably provides the most popular information from commercial genotyping companies such as *Ancestry.com* and *23andMe* to millions of customers. It also forms the basis of many standard population genetic analyses and most population genomic publications include ancestry inference analyses in one form or another (e.g., Rosenberg, et al. (2002); Li, et al. (2008)). Modern ancestry inference has roots in the seminal paper on STRUCTURE (Pritchard, et al., 2000). The model introduced in that paper assumes that each individual can trace its ancestry fractionally to a number of discrete populations. For each individual, independence is assumed between the two alleles at a locus, and the ancestry for each allele is then described as a mixture model in which the allele is assumed to be sampled from each of the ancestral populations with probability equal to the *admixture proportion* of this ancestral population. Many subsequent methods are based on the same model including FRAPPE (Tang, et al., 2005) and ADMIXTURE (Alexander, et al., 2009). Notice that this model implicitly assumes that the admixture proportions for each parent of an individual are the same. This assumption is arguably unrealistic for many human populations. In fact, for recently admixed populations, we would expect the admixture proportions to differ between the parents. However, the commonly used methods for admixture inference do not allow estimation of ancestry components separately for the two parents. We note that there is substantial information in genotypic data on parental admixture proportions. Even without linkage information, the genotypes can be used to infer parental ancestry. For example, consider the extreme case of a locus with two alleles, *T* and *t* at a frequency of 1 and 0, respectively, in ancestral population A, and a frequency of 0 and 1, respectively, in ancestral population B (i.e a fixed difference between two populations). Then the sampling probability of an offspring of genotype *Tt*, resulting from matings between individuals from populations A and B, is equal to one. However, if the two parents are both 50:50 (%) admixed between populations A and B, the probability, in the offspring, of genotype *Tt* is 0.5. In both cases the average admixture proportion of the offspring individual is 0.5. This is an extreme example, but it clearly illustrates that the offspring genotype distributions contain information regarding the parental genotypes that can be used to infer admixture proportions in the parents.

Recently, a method was developed for inferring admixture proportions, and admixture tracts, in the two parents separately from phased offspring genotype data (Zou, et al., 2015). This method models the ancestry process along each of the chromosomes as a semi-Markov process, as the length distribution of admixture tracts is well-known not to follow the exponential prediction of a Markov process (Gravel, 2012). It uses inference methods based on Markov Chain Monte Carlo (MCMC), Stochastic Expectation Maximization (EM), and a faster non-stochastic method for the case of a Markovian approximation to the ancestry process, and show that parental ancestry can be estimated with reasonable accuracy.

The objective of this paper is to explore the possibility of not only estimating admixture proportions in parents, but in grandparents, or even great grandparents. We show that the distribution of tract lengths provides information that can be used for such inference. By modeling the segregation of admixture tracts inside a pedigree we obtain a likelihood function that can be used to estimate admixture proportions in grandparents and great grandparents. While these estimates are associated with some variance, we show that they nonetheless can be used to distinguish between various hypotheses regarding the admixture proportions of parents, grandparents and great-grandparents. Our method has been implemented in a computer program called PedMix.

## Materials and methods

### Inferring admixture proportions from genetic data

We consider a single diploid individual from an admixed population. We assume two haplotypes *H*_1_ and *H*_2_ for this individual are given. Here, a haplotype is a binary vector of length *n*. *n* is the number of single nucleotide polymorphisms (SNPs) within the haplotype. Note that in real data *H*_1_ and *H*_2_ are usually inferred from the genotypes *G* and may have phasing errors. For the ease of exposition, we initially assume the absence of phasing errors in the haplotypes, and then extend the inference framework to allow phasing errors. The admixed population is assumed to be formed by an admixture of two ancestral populations (denoted as populations A and B) *g* generations ago. We assume *g* is known. For simplicity we assume there are two ancestral populations, although the method can be extended to allow more than two ancestral populations. We further assume allele frequencies in the two ancestral populations are known for all SNPs. Note that allele frequencies from extant populations that are closely related to the ancestral populations are typically available. For example, suppose the admixed individual has genetic ancestry in West Africa and Northern Europe. Then we may use the allele frequencies from the extant YRI and CEU populations, available from the 1000 Genomes Project (1000 Genomes Project Consortium, 2015), as approximations of the real ancestral allele frequencies. We also assume recombination fractions between every two consecutive SNPs are known. For human populations, recombination fractions are readily available (e.g. 1000 Genomes Project Consortium (2015)).

### Likelihood computation on the perfect pedigree model

The *perfect pedigree* model in Liang and Nielsen (2014) can be used to describe the segregation of admixture tracts. Here, an admixture tract is a segment of the genome which originates from a single ancestral population. This model differs from many of the models typically used for inferring admixture tracts of an extant individual (e.g. Tang, et al. (2006); Price, et al. (2009); Sankararaman, et al. (2008); Pásaniuc, et al. (2009)). This model directly models the segregation of admixture tracts within a pedigree. Most current models assume that the ancestry process follows a Markov chain along the chromosome. However, because of recombination between tracts from multiple ancestors, the exact process does not follow a first-order Markov process (Liang and Nielsen, 2014; Gravel, 2012). The perfect pedigree model establishes a more accurate, but also much more computationally demanding, model that does not assume a Markov process for the ancestral process, especially for recent admixture events.

Fig 1 illustrates the perfect pedigree model for an extant observed haplotype *H* at a single site. A perfect pedigree is a perfect binary tree where each node represents a haplotype. All internal nodes in the pedigree are ancestors of *H* (the single leaf in the pedigree). We trace the ancestry of *H* backwards in time until reaching the time of admixture, *T*_*m*_ generations ago. The 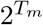 haplotypes at this time are called “founder” haplotypes (which themselves are unadmixed but may be from different ancestral populations). Under the assumption of no inbreeding, all ancestors are distinct. Notice that there is an assumption of a single admixture event. However, the model can easily be generalized to multiple admixture events.

**Figure 1:**
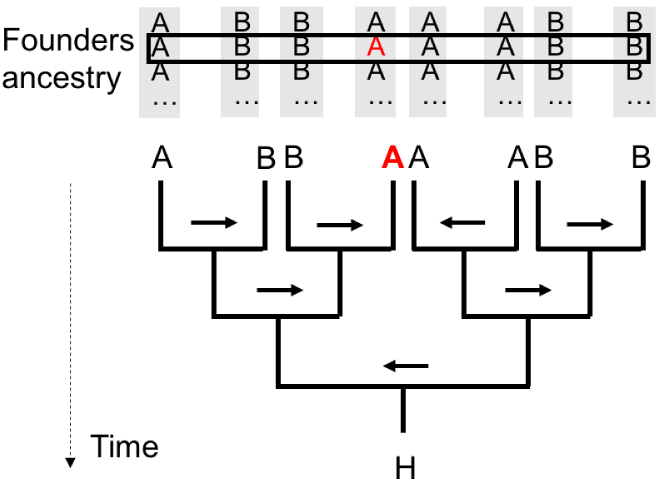
The perfect pedigree model for *T*_*m*_ = 3 for a single site. *H*: extant haplotype. Ancestry origins of 8 founders are listed above the perfect pedigree. At one site, A and B indicate which of the two ancestral populations of each founder’s haplotype. The combined vector of these values is *C*. Arrows: the recombination vector *R*. The population A shown in red: the ancestry of *H* as traced back by the recombination setting. Arrows can change direction at the next site.

There are two main aspects of the perfect pedigree: the ancestry vector *C* and the recombination vector *R*. *C* specifies which ancestral population each particular founder haplotype is from. For example, in Fig 1, *C* is a vector (*ABBAAABB*), of length 8. It indicates that the leftmost founder is from the ancestral population *A* while the rightmost founder is from ancestral population *B*. As founders are unadmixed, *C* does not change along the genome. *R* specifies from which of the two parental haplotypes each descendant haplotype inherits its DNA at a particular genomic position. *R* is the key component in the well-known Lander-Green algorithm (Lander and Green, 1987). As shown in Fig 1, one can visualize *R* as a set of arrows, one for each meiosis, pointing to the left or right. We have a list of recombination vectors for *n* sites (*R*_1_*, R*_2_, …, *R*_*n*_), where *R*_*i*_ is the recombination vector for site *i*.

The most obvious method for computing the likelihood *P* (*H M*) of the given haplotype *H* on the perfect pedigree model is using the Lander-Green algorithm (Lander and Green, 1987) to compute the probability of *H* for a given *C*. Then we sum these probabilities over all possible *C* to obtain *P* (*H|M*). Here *M* is a vector of admixture proportions for ancestors of interests in the pedigree. However, computation of *p*(*H|M*) directly using the Lander-Green algorithm is not practical for most datasets. This is because first we need to determine the ancestral setting, *C*, which specifies the ancestral population for each founder. Moreover, the number of possible *R* grows very fast with the number of generations in the pedigree. Note that the Lander-Green algorithm needs to enumerate all possible *R* values. Even considering just a single site *i*, there are 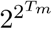 possible values of *C* and 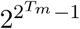 possible values of *R*_i_ (1 ≤ *i* ≤ *n*). These numbers are prohibitively large for e.g. *T*_*m*_ = 10. To circumvent this problem, we adopt a two-stage model as described below.

### A two-stage Markovian pedigree model for genotypes

Our objective is to infer the admixture proportions of ancestors in the perfect pedigree at the *K*^*th*^ generation in the past. Here, *K* is usually much smaller than the number of generations since admixture. For example, 1^*st*^ generation inference (*K* = 1) is for parents and 2^*nd*^ generation inference (*K* = 2) is for grandparents. The first phase of the two-stage model involves modeling the first *K* generations in the past using the perfect pedigree model. In the second phase, starting at the *K*^*th*^ generation in the past, there are 2^*K*^ ancestors, which are assumed to have ancestry distributions following the standard Markovian model. The ancestry of these 2^*K*^ founders can change along the genome following the standard Markovian process. This allows us to model the admixture of recent ancestors (e.g. parents and grandparents) without explicitly considering the entire pedigree.

The model defined so far concerns haploid genomes/chromosomes. However, most real data are from diploid individuals, possibly with unknown or relatively poorly estimated haplotype phasing. We extend the two-stage pedigree model by assuming that each of the two haplotype from the extant individual has been estimated, but with phasing errors that occur at a constant switch error rate. This leads to a genotype-based perfect pedigree model.

Fig 2 (A) illustrates the genotype-based perfect pedigree at a single position. It consists of two perfect pedigrees, one for each of the two haplotypes *H*_1_ and *H*_2_. Each node in the outline tree denotes an ancestral genotype of the extant genotype *G*. The two haplotypes *H*_1_ and *H*_2_ of *G* follow different pedigrees independently. For simplicity, we use a single haplotype with “average” admixture tracts to represent a diploid founder, which works well in practice. Note that the estimated admixture proportion of a founder is the average of the admixture proportions of its two haplotypes. One can view this “average” haplotype has the admixture proportion equal to the diploid founder.

**Figure 2:**
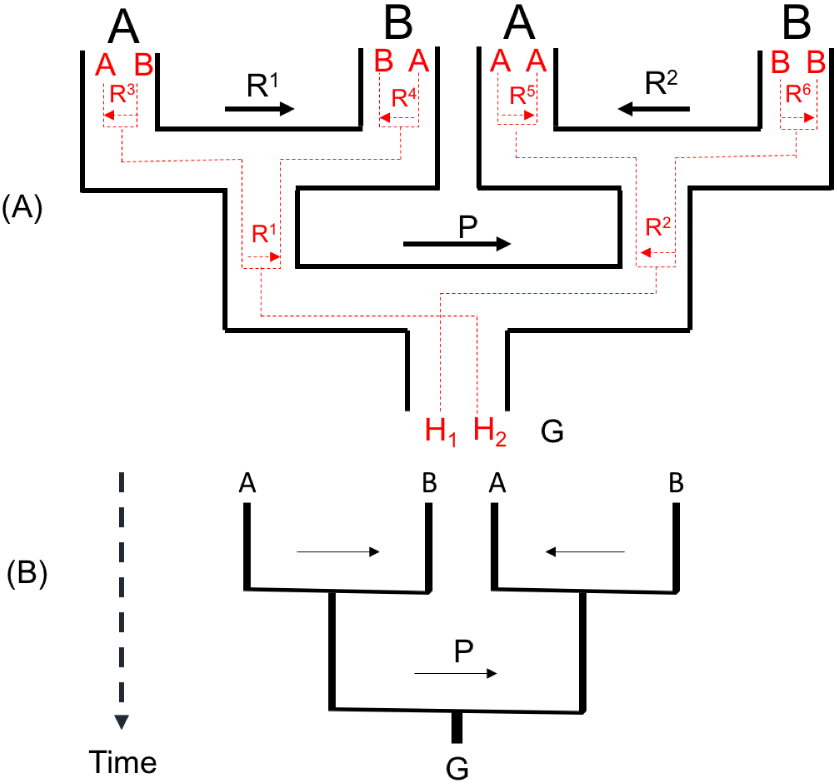
The perfect pedigree model for genotype *G* = (*H*_1_, *H*_2_) at a site, and *K* = 2. (A) Outline pedigree in black: the perfect pedigree for genotype *G*. Two pedigrees embedded in red are for haplotype *H*_1_ and *H*_2_ respectively. Ancestral settings and recombination settings with the same label have the same meaning. (B) The simplified perfect pedigree for genotype *G*. Ancestral vector *C*: (*ABAB*). The arrows without label define recombination vector *R*. *P*: the phasing error setting.

To allow phasing errors between *H*_1_ and *H*_2_, we introduce the phase-switching indicator *P*. It indicates whether at this position the two haplotypes switch or not. One can visualize *P* as the arrow labeled by *P* in Fig 2. A *P*-arrow pointing to the left indicates that *H*_1_ traces to the left half of the pedigree and *H*_2_ traces to the right half of the pedigree. A *P*-arrow pointing to the right indicates the opposite. When moving along the diploid sequence (genotype), the direction of *P* changes when a phasing error occurs. Thus we can combine the two pedigrees for *H*_1_ and *H*_2_ and let the two haplotypes from a single individual collapse into one node, as illustrated in Fig 2 (B).

The full information regarding the ancestry of a genotype, *G* = {*H*_1_, *H*_2_}, in a fixed pedigree is then given by the ancestral configuration *AC* = (*P, C, R*). The sampling probability of *G* can be computed naively by summing over all possible *AC*s. The ancestral configuration *AC* naturally leads to an Hidden Markov Model (HMM) that can be used for efficient calculation of the likelihood.

Let 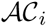 denote a set containing all possible ancestral configurations at site *i* and *AC*_*i*_ denote an element that belongs to 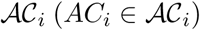. In a perfect pedigree of *K* generations, *AC*_*i*_ = (*P*_*i*_, *C*_i_, *R*_*i*_) is a binary vector of 2^*K*+1^ − 1 bits and represents a state at the site *i*. For each state, *P*_*i*_ has exactly one bit where a “0” (respectively “1”) represents the phasing arrow pointing to the left (respectively right) and “1” represents the phasing arrow pointing to the right. *R*_*i*_ is a binary vector of 2^*K*^ − 2 bits indicating the recombination states associated with all 2^*K*^ − 2 meiosis in the pedigree, where “0” (respectively “1”) represents a recombination arrow pointing to the left (respectively right). *C*_*i*_ is a binary vector that indicates the ancestry of each of the 2^*K*^ ancestors and contains 2^*K*^ bits when there are two ancestral populations. Also, if *C*[*j*] = 0 (respectively *C*[*j*] = 1) the *j*-th founder is from the population *A* (respectively *B*) at the current site. In the example in Fig 2(B), *AC* = (*P, C, R*) at this site can be expressed as the binary vector (1, 0101, 10).

We define *h*(*AC*_*i*_) as the joint probability of the length-*i* prefix of *G* (i.e. *G*[1*..i*]) and the ancestral configuration *AC*_*i*_ at the site *i*. Given a genotype *G* with *n* sites, the likelihood 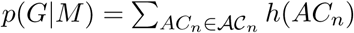. The critical step is the computation of *h*(*AC*_*i*_) for each configuration *AC*_*i*_ at the site *i*. This can be carried out in a recurrence for *i* ≥ 2:

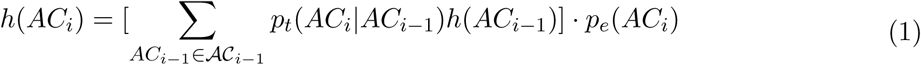

where *p*_*t*_(*AC*_*i*_|*AC*_*i*−1_) is the transition probability from *AC*_*i*−1_ at the site *i −* 1 to *AC*_*i*_ at the site *i* and *p*_*e*_(*AC*_*i*_) is the emission probability of an allele given the ancestral configuration *AC*_*i*_ at the site *i*. This is the standard forward algorithm for HMMs. Details are given below. Transitions in the HMM may occur between adjacent sites and we assume, for generality, that the configurations at sites *i* − 1 and *i* are fully connected as illustrated in Fig 3.

#### Transition and Emission probabilities of the HMM

Consider a founder *j* and two sites that are separated by *d* nucleotides. We first define the one-step ancestry transition probabilities 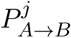 and 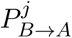. 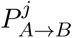 (respectively 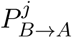) is the probability that the ancestral population *A* (respectively *B*) changes to ancestral population *B* (respectively *A*) along the genome for the founder *j* when *d* = 1. Recall that the ancestry process of an ancestor follows a standard Markovian model. Suppose a haplotype of the individual *j* has the ancestral population *A* at the site *i* − 1. The probability that the site *i* has the ancestral population *B* is approximately 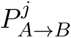 and the probability that the site *i* has the ancestral population *A* is approximately 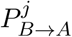, assuming that *d* (the number of bp between the sites *i* and *i* − 1) is small. Multiple transitions in the interval are ignored. We then define *T*_*j*_ to be the *d*-step transition probability of the ancestral settings for ancestor *j*:

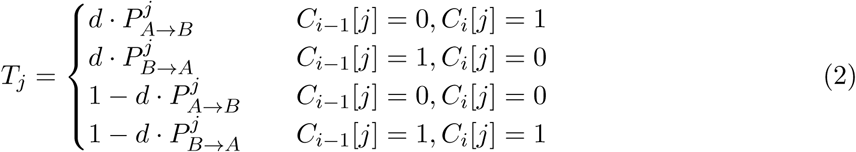

**Figure 3:**
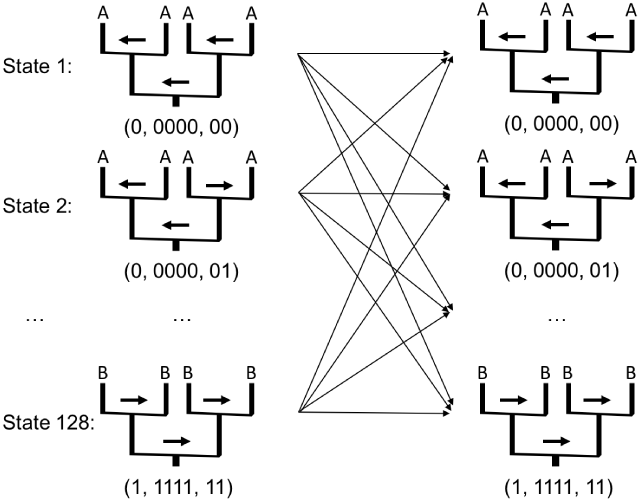
An example of 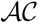-based HMM with 128 *ACs* as states for *K* = 2 generations (i.e. grandparents) with arrows indicating possible transitions along the Markov chain from site *i* − 1 to *i*. The vectors under each pedigree provide the binary representations of *P*, *C*, and *R*, respectively, for the pedigree. The two top thick arrows and the lower thick arrow indicate the settings of *R* and *P*, respectively.

Notice that this is a function of *d*. *i*, and *i* − 1 are suppressed in the notation. Using similarly simplified notation, we define the phasing transition probabilities *I* as

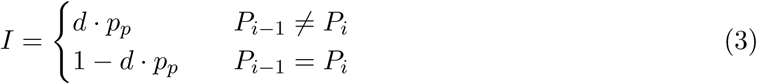

where *p*_*p*_ is the probability of a phasing error per unit length (assumed to be known and small enough that double or more phasing errors can be ignored).

We also define *B*_*k*_ as the transition probability of the recombination vector for the *k*th bit. Given the recombination map of *G*, the recombination probability *B*_*k*_ between the two sites is computable. Let 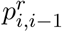 denote the probability of one recombination event between sites *i* and *i* − 1, then

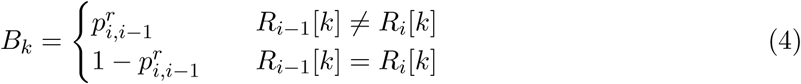

Using this simplified notation, and assuming independence among transitions associated with recombination, phasing errors and the ancestral population setting, the transition probabilities of the Markov chain are then given by:

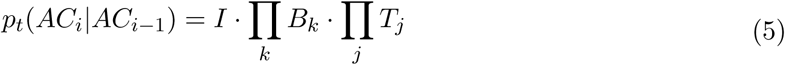

As mentioned above, the emission probability at the site *i* is a function of the ancestral population assignment, *C*_*i*_, and the alleles of the focal individual. At the site *i* of the genotype *G*, there are two haplotypes (*h*_1_*, h*_2_). Let 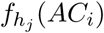 be the allele frequency in the population specified by *AC*_*i*_ for the allele observed at the position *i* of *h*_*j*_ (*j* = 1, 2). The emission probability is then

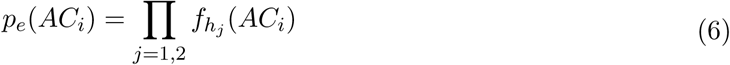

as in the standard definitions in genetic ancestry models (e.g., Pritchard, et al. (2000)).

#### Fast computation of sampling probability in PedMix

The main computational burden in the evaluation of Equation 1 is that the calculation of *h*(*AC*_*i*_) requires a multiplication of the transition probability matrix and the vector *h*(*AC*_*i*−1_). This leads to a computational complexity of 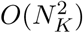, where *N*_*K*_ is the number of possible states in 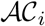. This is a significant burden on computation: for example if *K* = 3, the run time is on the order of 2^30^. To address this problem, we have developed a divide and conquer algorithm for computing the probability of *ACs*, which runs in *O*(*N*_*K*_*log*(*N*_*K*_)) time.

Let *P*_*i*_ denote the probability vector that contains all *h*(*AC*_*i*_) for 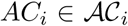 at the site *i*. Let *T*_*i*−1,*i*_ denote the transition probability matrix containing the transition probabilities *p*_*t*_(*AC*_*i*_|*AC*_*i*−1_) that one AC at the site *i* − 1 transits to another AC at the site *i*. To obtain *P*_*i*_, we need to compute *T*_*i*−1,*i*_*P*_*i*−1_. Direct computation leads to quadratic complexity.

For simplicity, we omit the site index notation *i* or *i* − 1 in *T*_*i*−1,*i*_ and *P*_*i*−1_. Let *T*^*b*^ denote the transition probability matrix for *AC* that has *b* bits. The *AC* is represented as a binary vector of length *b*. Let *P*^*b*^ denote the probability vector for the previous site (*i* − 1) of length *b*. A bipartition of a matrix is a bipartition of each dimension, which divides a matrix into four sub-matrices with equal size. A bipartition of a vector is a division that equally cuts the vector into two sub-vectors. Fig 4 shows an example of a transition probability matrix *T*^3^ and a probability vector *P*^3^ for *AC* with 3 bits. For example, the (2, 3) element in *T*^3^ is the transition probability *p*_*t*_((001) (010)). The bipartition for *T*^3^ and *P*^3^ is shown as red lines.

**Figure 4:**
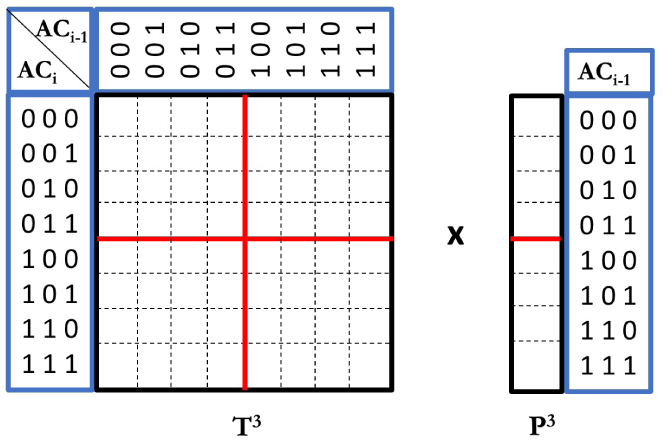
Faster calculation of the probabilities of ACs. Red lines break the transition probability matrix into four smaller pieces (for ACs with length 2). The probability vector at the previous site is broken into two pieces. Multiplication of the matrix and the vector is faster due to shared parts between these pieces.

We observe that each bit in an *AC*_*i*−1_ transits to a bit in *AC*_*i*_ independently (i.e. the transition probability of each bit in *AC*_*i*_ doesn’t depend on other bits). We use 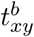 to denote the transition probability of the *b*th bit from *x* to *y* (*x, y* ∈ {0, 1}). We can adapt the divide-and-conquer approach in Idury and Elston (1997) to our problem as follows. With bipartition, *T*^*b*^ can be viewed as four sub-matrices, and *P*^*b*^ can be divided into two sub-vectors. The key of the divide and conquer approach is given in the Eq 7.

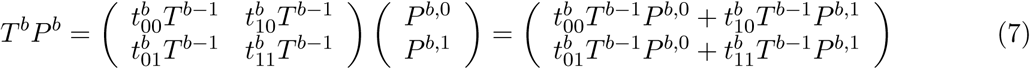

Each sub-matrix of *T*^*b*^ is equal to *T*^*b*−1^ multiplied by 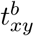. Here, *T*^*b*−1^ is a transition probability matrix where the *b*th bit of *T*^*b*^ is masked off. For example, the top left sub-matrix of *T*^3^ (Fig 4) is equal to 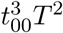 and the top right sub-matrix is equal to 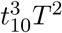. Let *P*^*b*^ = (*P*^*b*,0^, *P*^*b*,1^) denote the bipartition of the probability vector. Then *T*^*b*^*P*^*b*^ can be computed by computing *T*^*b*−1^*P*^*b*,0^ and *T*^*b*−1^*P*^*b*,1^. In general, *T*^*b*−1^*P*^*b*,0^ and *T*^*b*−1^*P*^*b*,1^ can then be divided in a similar way until we reach *T*^1^ (masking off *b* − 1 bits in *AC*). For the *K*^*th*^ generation inference, each *AC* has *b* = 2^*K*+1^ − 1 bits, which leads to *N*_*K*_ = 2^*B*^ = 2^2^ −^1^ possible states at each site. The divide and conquer scheme reduces computational complexity from 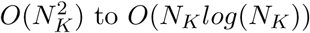 to *O*(*N*_*K*_*log*(*N*_*K*_)).

### Probabilistic inference

Maximum Likelihood (ML) inference of admixture proportions can be obtained by maximizing the sampling probability *p*(*G|M*) of the 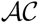-based HMM model:

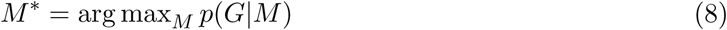

Let 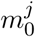 and 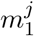 denote the admixture proportions of the populations *A* and *B* respectively for the ancestor *j*. These admixture proportions are then given by the stationary frequencies of the Markov chain, which according to standard theory are given by

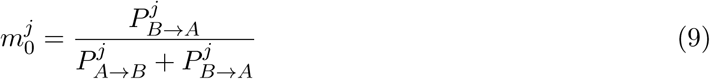

and

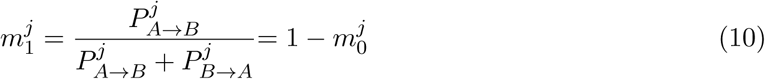

respectively. From the invariance principle of ML, it follows that if 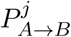 and 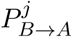 are estimated by ML, the resulting estimates of 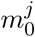 and 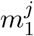 are also ML estimates.

To obtain ML estimates of 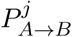 and 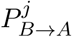 we apply the Boyden-Fletcher-Goldfarb-Shanno (BFGS) method of optimization. We use an implementation of the limited-memory version of the algorithm (L-BFGS) (http://www.chokkan.org/software/liblbfgs) and the finite difference method for estimating derivatives. We transform bounded parameters using the logit function to accommodate bound constraints.

### Preprocessing

There are several aspects of real data that are not considered by our models and may affect the inference accuracy, in particular background Linkage Disequilibrium (LD) and phasing errors. Background LD refers to non-random association between alleles not caused by admixture. Background LD may mislead HMM methods which assume conditional independence among SNPs. As a consequence it may confuse the background LD with the admixture LD. Phasing errors may also introduce an extra layer of noise.

The traditional approach for addressing the problem of background LD is to trim the data sets by removing SNPs. We compare two possible strategies for doing this:

1. Data trimming based on allele frequency differences (frequency-based pruning).
2. Data trimming based on LD patterns (LD pruning).

Frequency-based pruning relies on a trimming threshold *d*_*f*_, which specifies the minimum allele frequency difference in the two source populations. A SNP site is trimmed if the absolute difference between the allele frequencies in the source populations is smaller than *d*_*f*_. See the Supplemental Materials for more details on this approach.

In LD pruning, SNPs are removed in order to minimize the LD among SNPs located in the same region. This is the more commonly used strategy implemented in programs such as PLINK (Purcell, et al., 2007). See the Supplemental Materials for more details on LD pruning.

The advantage of the second approach is that it more directly reduces LD in the data. The advantage of the first approach is that it keeps the most ancestry informative SNPs in the data set. Both approaches improve inference accuracy and reduce computational time. However, our implementation of frequency-based pruning leads to slightly better performance and we, therefore, use this method as the default unless otherwise stated.

### Phasing error

In real haplotype datasets, phasing error usually cannot be eradicated when haplotypes are inferred from genotypes. In some sense, phasing errors and recombination have similar effects on the genomes of the extant individual. We have developed a technique for removing some phasing errors during preprocessing. Briefly, we first estimate the admixture tracts for the current haplotypes. We expect admixture tracts to be relatively long, but may be shortened by Phasing errors. Phasing errors can, therefore, be removed to some extend by removing unexpectedly short admixture tracts. See the Supplemental Materials for details.

## Results

This section contains results on simulated, semi-simulated and real data. Some results are given in the Supplemental Materials.

### Results on simulated data

#### Simulation settings and evaluation

We perform extensive simulations to evaluate the performance of our method. We first simulate a number of haplotypes using macs (Chen, et al., 2009) from two ancestral populations which diverged from one ancestral population at 4*N*_*e*_*t* generations in the past. Here *N*_*e*_ is the effective population size. An admixed population is then formed by merging the two ancestral populations and simulating the process of random mating, genetic drift, and recombination using a diploid Wright-Fisher model for *g* additional generations. We model recombination rate variation using the local recombination estimates from the 1000 Genomes Project (1000 Genomes Project Consortium, 2015). The hotspot maps of the 22 human autosomal chromosomes are concatenated for a single string of 3 × 10^9^*bp* and subsequently simulated genomes are divided into 22 chromosomes of equal length to facilitate clearly interpretable explorations of the relationship between accuracy and the amount of data. Haplotypes are paired into genotypes and phasing errors are then added stochastically, by placing them on the chromosome according to a Poisson process with rate *p*_*p*_. By default, no phasing error is included in the simulations. The parameters we use in the simulations are listed and explained in Table 1 together with their default values. For the default setting, the approximate total number of SNPs simulated by macs is ~ 14.7*M*. Here we apply frequency-based pruning to trim data (see Results section). Frequency-based pruning removes SNPs with a minor allele frequency difference in two ancestral populations less than the pruning threshold *d*_*f*_. After pruning with the default *d*_*f*_, each of the 22 chromosomes contains ∼ 26, 000 SNPs. In some cases, the default simulated length *L* = 3 × 10^9^*bp* results in high computational burden. Therefore, in some simulations we also use a shorter length of *L* = 5 × 10^8^*bp*, divided into 3 chromosomes. If not otherwise stated, we use *L* = 3 × 10^9^*bp* to be the default setting.

**Table 1:** A list of parameters and their default values used in the simulation

| Description | Symbol | Default |
| --- | --- | --- |
| The number of haplotypes | $n_h$ | 1000 |
| The number of chromosomes | $n_c$ | 22 |
| Effective population size | $N_e$ | 10000 |
| Region length (bp) | $L$ | $3 \times 10^9 / 5 \times 10^8$ |
| Mutation rate (per generation per bp) | $\mu$ | $1 \times 10^{-8}$ |
| Recombination rate (per generation per bp) | $\rho$ | $1 \times 10^{-8}$ |
| Ancestral populations splitting time (scaled with $N_e$) | $t$ | 0.2 |
| The number of generations for admixture | $g$ | 10 |
| The number of individuals to infer | $n_i$ | 10 |
| Frequency-based pruning threshold | $d_f$ | 0.5 |

PedMix is then applied to the simulated genotype data from the admixed population for inference of admixture proportions of ancestors in the 1st generation (parents), the 2nd generation (grandparents) and so on. To evaluate accuracy, we use the mean absolute error (MAE) between the estimated admixture proportion, 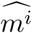, and the true admixture proportion, *m*^*i*^, for the *i*th ancestor in the *K*^*th*^ generation, as the metric of estimation error. If there are multiple individuals, we further take the average over all individuals, i.e. the mean error for *n* individuals is defined by Equation 11. Without loss of generality, we only consider the estimate of the proportions of the ancestral population A. Because we assume two ancestral populations, the expected mean errors of admixture proportions for two ancestral populations are identical.

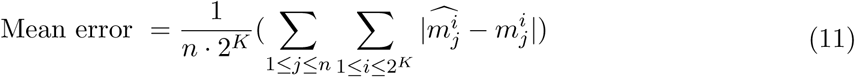

As the admixture proportions inferred by the method are unlabeled with respect to individuals, this leads to ambiguity on how to match the inferred proportions to the true proportions. We address this problem using a “best-match” procedure by rotating the parents for each internal node in pedigree to find the best match between the inferred and the simulated admixture proportions. For example, for inference in parents, we have true admixture proportions for two parents (*m*^1^*, m*^2^) and estimated admixture proportions 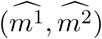. We match both 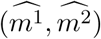 and 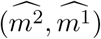 to (*m*^1^, *m*^2^) and choose the one with smaller mean error. For the case of grandparent inference, there are eight possible matchings and we explore all eight to obtain the best match.

#### Evaluation of ancestral inference accuracy

Figure 5 shows the mean error when inferring admixture proportions of parents, grandparents, and great grandparents under the default simulation settings (as shown in Table 1). We compare the performance of PedMix to what is expected from random guess based on a Bayesian model. The random guess is described in the Supplemental Materials. We sample 10 genotypes from simulated admixed population for *g* (*g* ≥ 3) generations.

**Figure 5:**
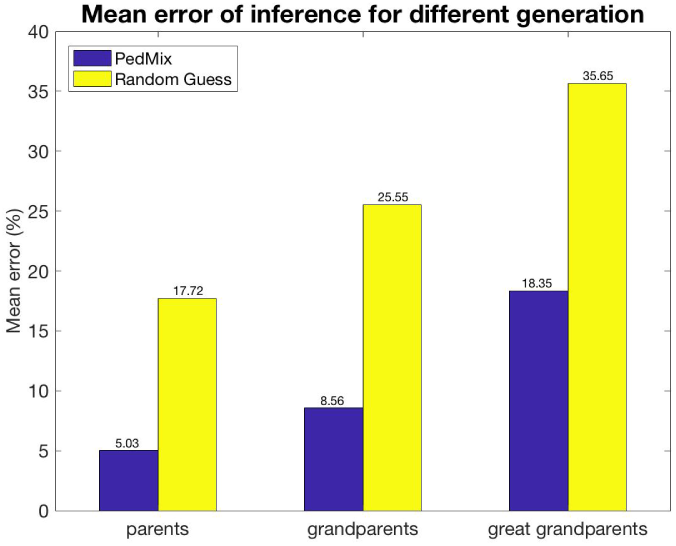
Comparison between the accuracy of PedMix and a random guess for parents, grandparents and great grandparents. About 570,000 SNPs are used for parent and grandparent simulations. About 26,000 SNPs are used for great grandparent simulations since this case needs more computing resources.

Note that inference of great grandparents admixture proportions is computationally demanding in the current framework. Therefore, we use a more extreme trimming threshold, *d*_*f*_ = 0.9, when inferring great grandparent admixture, resulting in only 26,638 SNPs.

As expected, it is easier to estimate admixture proportions of more recent ancestors. This is because, as we trace the ancestry of a single individual back in time, the genome of the extant individual contains progressively less information about an ancestor.

#### Comparison of PedMix to existing methods

Although there are no existing methods for inferring the admixture proportions of grandparents and great grandparents that we can compare PedMix to, there is a method called ANCESTOR (Zou, et al., 2015) that infers admixture proportions of parental genomic ancestries given ancestry of a focal individual. And there are many methods (e.g., ADMIXTURE (Alexander, et al., 2009) and RFmix (Maples, et al., 2013)) for inferring admixture proportions of individuals of the current generation. In this section we first compare estimates of the admixture proportions of a focal individual obtained from ADMIXTURE and RFmix, arguably the state-of-the-art methods for ancestry inference, to the average of parental or grandparental admixture proportions inferred using PedMix. Here, we use the average of the estimated admixture proportions from ancestors as the proxy for the admixture proportion of the focal individual. If the admixture proportions of ancestors inferred by PedMix are accurate, we would expect the average of these admixture proportions of ancestors can serve as a good approximation for the focal individual. And this average should be approximately as accurate as the admixture proportions inferred by RFmix and ADMIXTURE. This is verified with simulation data in the Supplemental Materials. We note that the high accuracy of PedMix in inferring the admixture proportion of a focal individual from the average of parental or grandparental proportions does not necessarily imply that the parental and grandparental admixture proportions themselves are accurately inferred. However, if the admixture proportion of a focal individual is poorly estimated from the inferred admixture proportions of the ancestors, this may suggest that the admixture proportion estimates for the ancestors also are not accurate.

We randomly sample 20 individuals from an admixed population and run ADMIXTURE, RFmix and PedMix on the same datasets. The genotypes are preprocessed with LD pruning (see the Supplemental Materials) and contain phasing errors simulated with rate *p*_*p*_ = 0.00002 per *bp*. We deduce the ancestry of each individual using PedMix from the inferred admixture proportions of either parents or grandparents, by using the average of the inferred admixture proportions of the ancestors. ADMIXTURE and RFmix infer the admixture proportions of extant individuals directly. More details on how ADMIXTURE and RFmix are applied are given in the Supplemental Materials. Table 2 shows the mean error as defined in Equation 11 and the error’s standard deviation. Our results show that the admixture proportions inferred from the average of ancestral admixture proportions in PedMix are comparable to those of RFmix and ADMIXTURE. The estimate by parents matches the results of RFmix and ADMIXTURE. The estimate by RFmix and ADMIXTURE is slightly better than the estimate by grandparents.

**Table 2:** **Mean and standard deviation** of the error in the estimate of admixture proportions for ADMIXTURE, RFmix and PedMix (in unit of %). Note the PedMix results are the average proportions from the estimated admixture proportions of parents (denoted as par.) or grandparents (denoted as grandpar.).

| Error (in %) | ADMIXTURE | RFmix | PedMix (par.) | PedMix (grandpar.) |
| --- | --- | --- | --- | --- |
| mean | 1.16 | 1.77 | 2.01 | 3.53 |
| standard deviation | 1.43 | 1.52 | 2.5 | 2.53 |

We further compare the estimates of admixture proportions of parents from PedMix to those from ANCESTOR. ANCESTOR requires that the ancestry states and the tract lengths be provided as input for the algorithm. To accomplish this, we use the ancestry tracts inferred by RFmix when running ANCESTOR. More details on how ANCESTOR is applied are given in the Supplemental Materials. Mean error is computed between the true admixture proportions of parents and the estimates from ANCESTOR or PedMix (Table 3). Estimates by PedMix are substantially more accurate than ANCESTOR (approx 6.5% error versus 9.6% error).

**Table 3:** **Mean and standard deviation** of the error in the estimate of admixture proportions of parents from ANCESTOR and PedMix (in unit of %).

| Error (in %) | ANCESTOR (parents) | PedMix (parents) |
| --- | --- | --- |
| mean | 9.63 | 6.54 |
| standard deviation | 6.13 | 2.14 |

#### Impact of simulation parameters

We perform additional simulations to investigate the impact of various simulation parameters on the accuracy of our method. To investigate the effect of mutation rates and recombination rates, we use the default setting with a shorter genome of length *L* = 5 × 10^8^ (Table 1) to reduce the computational time (Fig 6). The expected number of SNPs simulated in a region increases linearly with the mutation rate. This leads to a reduction in the mean error with increased mutation rates, as more informative markers are available for analysis (Fig 6 (A)). However, the reduction is modest because the statistical accuracy is mostly limited by the number of admixture tracts and not by the number of markers. In contrast, recombination rate has a much stronger effect on the accuracy than mutation rate because increased recombination rates introduce more admixture tracts (Fig 6 (B), and also Supplemental Materials). The mean error for both parental and grandparental inferences decreases and then asymptotes as recombination rate increases further. When the recombination rate increases to more than 5 × 10−^8^, the improvement in accuracy becomes smaller, especially for parental inference (also see the Supplemental Materials). As the length of each tract decreases, the information regarding the ancestry for each tract also decreases. Even with very high recombination rates, there may still be some error determined by the degree of genetic divergence between populations and the number of generations since admixture.

**Figure 6:**
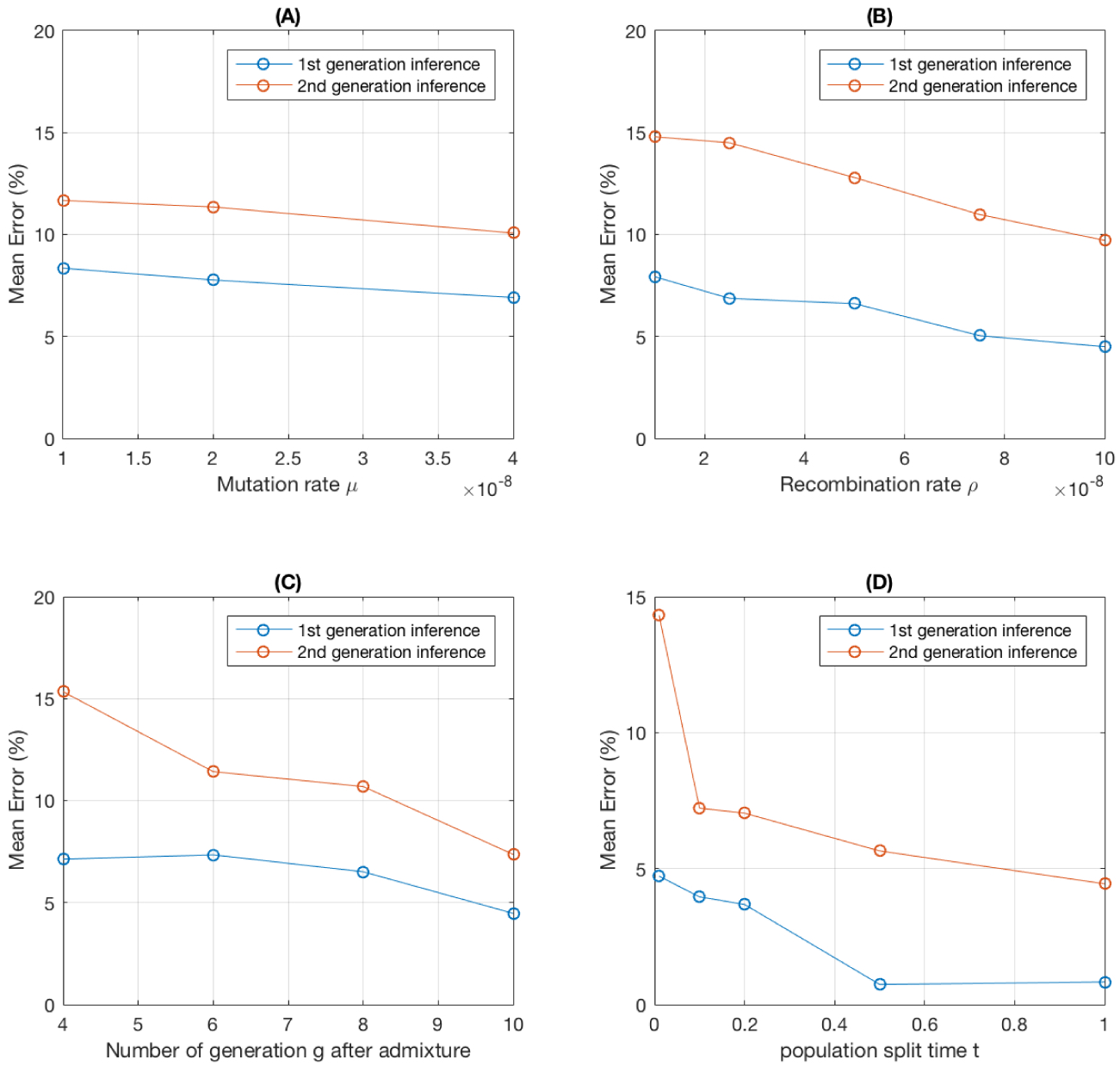
Mean error for different simulation parameter settings. (A) Varying mutation rates. *L* = 5 × 10^8^. (B) Varying recombination rates. *L* = 5 × 10^8^. (C) Varying the time since admixture. (D) Ancestral population split time. For *t* = 0.01, there are no SNPs left after using the *d*_*f*_ = 0.5 cut-off. As a result, we use *d*_*f*_ = 0.2, leaving 74,227 SNPs for analysis. Default parameters are used except for the variable indicated by the X axis of each plot.

The simulations assume a model of two ancestral populations that diverged 4*N*_*e*_*t* generations ago and then admixed *g* generations ago. The performance of the method clearly depends on these parameters. If *g* is small, the number of admixture tracts is also small, complicating inferences, particularly in the grandparental generation. As *g* increases from 4 to 10, the mean error reduces from 7.13% to 4.47% for parent inference and from 15.34% to 7.36% for grandparents respectively (Fig 6 (C)). There is also a strong effect of *t* on the accuracy. As *t* increases, allele frequency differences between the admixing populations increase and it becomes easier to distinguish admixture tracts from two ancestral populations (Fig 6 (D)). When *t* > 0.5 the mean error for parental inferences drops to below 1%.

#### Phasing error

Real data may contain phasing error. We have implemented a preprocessing approach for reducing the phasing error. See the Supplemental Materials for details. To evaluate the effect of phasing error, we simulate data with the phasing error rate 2 × 10−^5^ per bp. We then compare the mean error by PedMix using genotype data without phasing errors, genotype data with phasing errors, and genotype data with phasing errors preprocessed to remove some phasing errors. As for the data generated with phasing error rate 2 × 10−^5^, we run PedMix directly without preprocessing. Then we use the technique described in the Supplemental Materials to preprocess the data.

As shown in Figure 7 (A), phasing error reduction by preprocessing increases the inference accuracy. Note that at the default setting, the phasing error rate is nearly 2,000 times larger than the recombination rate, which can affect the accuracy of the method significantly. Thus, in real data analyses, it is important to reduce the phasing error in some way.

**Figure 7:**
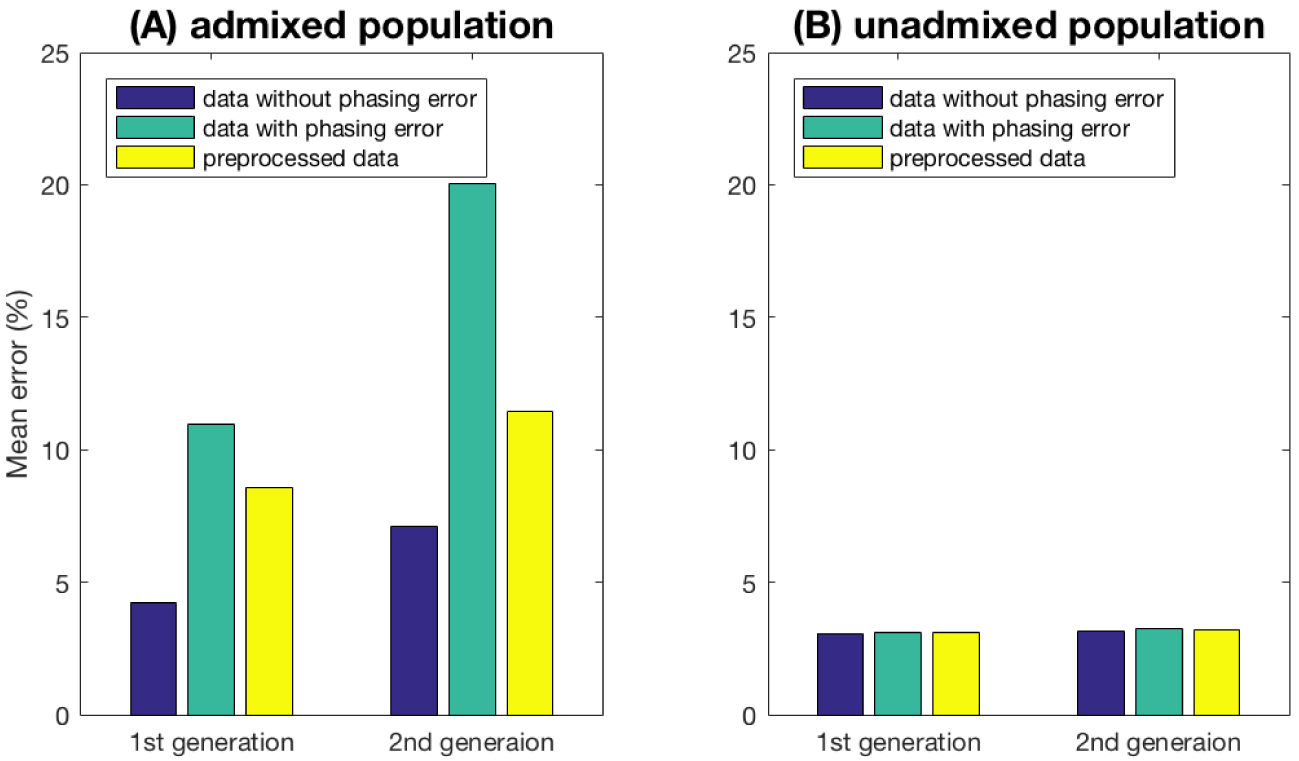
Inference error vs phasing error: comparison among three different datasets: dataset without phasing errors, dataset with phasing errors and preprocessed datasets. (A) Samples from an admixed population. (B) Samples from an unadmixed population.

We also consider the case of unadmixed individuals. In that case, phasing errors may be interpreted as recombination events during inference, but the overall admixture proportion estimates should be relatively unaffected. To illustrate this point, we sample 10 individuals from the same population and run PedMix to infer their admixture proportions. The performance of PedMix is very stable as shown in Figure 7 (B), with an inference error of approximately 3% for parents and grandparents.

Phasing error adds noise to the model, especially in the region where the two haplotypes have different ancestral states. As we decrease the effect of the phasing error in the data using preprocessing, the inference error decreases significantly.

### Results on semi-simulated data

We now show results on semi-simulated data. Here, we use genotypes of CEU/YRI/ASW populations from the 1000 Genomes Project as the founders of a fixed pedigree topology as shown in Figure 8. This way, the genotypes are closer to the real data and we know the origin of the founders. For this pedigree of two generations, we select four genotypes from one or more populations among CEU, YRI and ASW populations as grandparents. We assume there is no phasing error along these grandparental genomes. Then we simulate two genotypes as parents and one genotype as the focal individual based on the standard genetics law. Recombination rate is modeled from the hotspot maps of the 1000 Genomes Project as in the other simulations. To assess the impact of phasing errors, we also create data with phasing errors by adding phasing errors stochastically with the rate *p*_*p*_ = 0.00002 per *bp* for the focal individuals. We run PedMix on the genotype of the focal individual genotype with or without phasing error to infer the admixture proportions of parents and grandparents. RFmix is run to estimate the admixture proportions of parents (respectively grandparents) using the genotypes of parents (respectively grandparents). We use the estimates from RFmix with the ancestral genotypes as the ground truth on the admixture proportions of ancestors. Here we examine six cases with different ancestral origins of the grandparents: CCCY, CCYY, CYCY, AAAA, AAAC and AACY (where C is for CEU, Y is for YRI and A is for ASW). As an example, CCCY stands for the four grandparents from CEU, CEU, CEU and YRI respectively. Figure 8 shows the estimates by RFmix and PedMix. Mean error is computed from the six inferred admixture proportions (two parents and four grandparents) in the pedigree and their estimates by RFmix. Although the focal individuals in the pedigrees CCYY and CYCY both have around 50% admixture proportion, PedMix is able to tell the difference in the parents by estimating the parental admixture proportions being 82.41% and 16.43% for CCYY and 47.05% and 43.78% for CYCY. This largely agrees with the true admixture proportions of the parents, which are 99.91% and 0.09% for CCYY and 50.04% and 49.96% for CYCY. Note that the true admixture proportions for the two parents are known for a pedigree with the known grandparental origin. For example, in the CCYY case the true parental admixture proportions are 99.91% and 0.09%. This is because one parent has two CEU grandparents and thus this parent is about 100% CEU. Similarly, the other parent is around 100% YRI. This indicates that PedMix is able to collect useful information from the admixture tract lengths in the focal individual. Results on genotypes without phasing errors tend to be more accurate than those with phasing errors. Our results indicate that phasing errors can indeed lead to larger inference error for some cases. Thus, it is useful to use haplotypes with less phasing errors. Estimates for parents are more accurate than those of grandparents.

**Figure 8:**
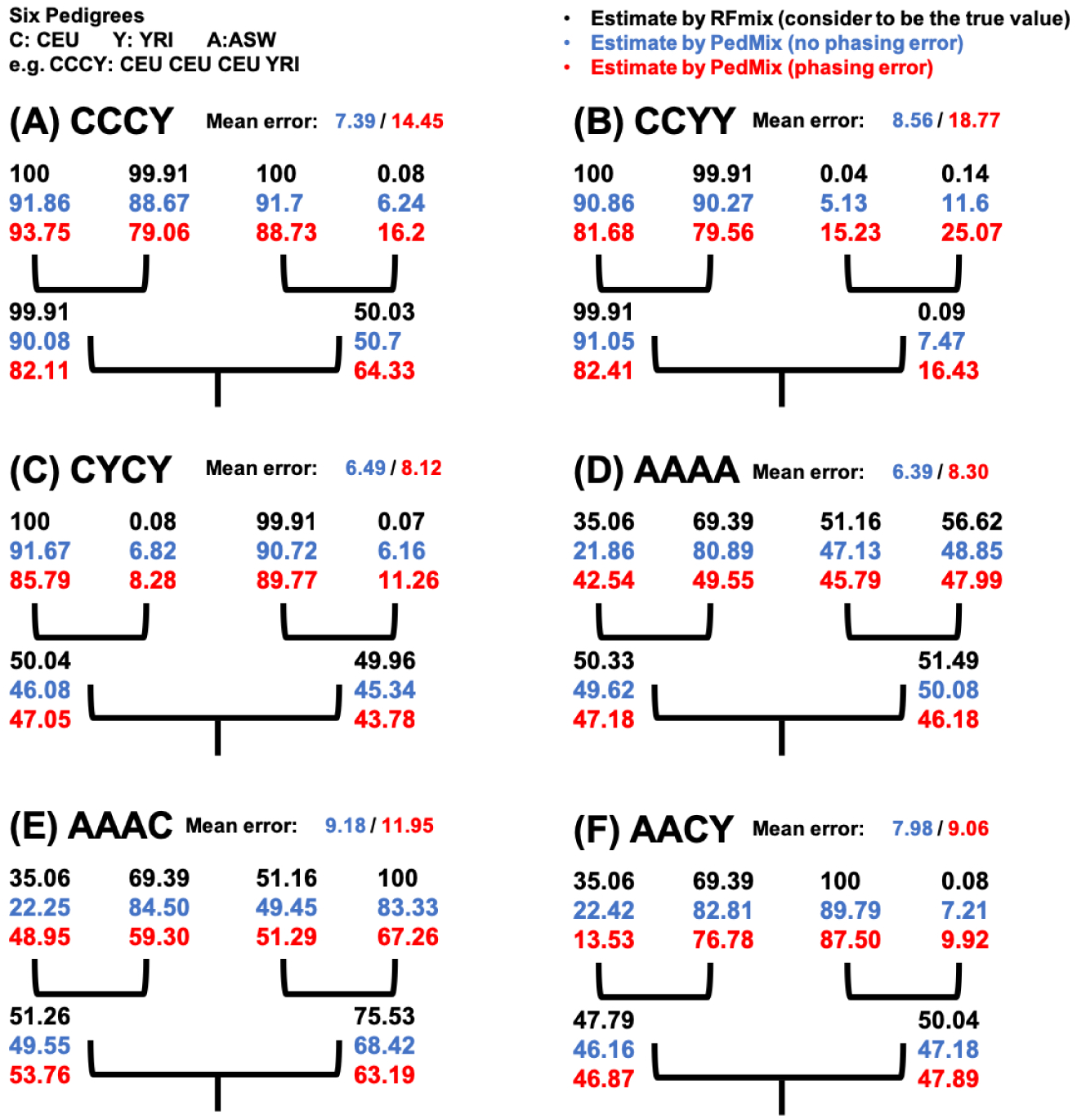
Six pedigrees used for the simulations scheme of semi-simulated data. The percentage of CEU origin is shown in the pedigrees. Admixture proportions in black are estimates by RFmix using the ancestors’ genotypes directly and are assumed to be the true proportions. Admixture proportions in blue and red are estimates by PedMix using genotypes of the focal descendant individual with (red) or without (blue) phasing error. Mean error is the average over the two parents and four grandparents.

### Results on real data

To evaluate the performance of PedMix on real genetic data, we run PedMix on the haplotypes of ten trios from the ASW population from the HapMap data (International HapMap 3 Consortium, 2010). Note that the exact ancestry of parents and grandparents in these trios are not known. Here, for parents, we use the inferred admixture proportions by running RFmix on the given haplotypes of parents as the true admixture proportions of the parents. The case of grandparents is more difficult because genetic data of grandparents is not available from the HapMap project. In order to evaluate the performance on grandparents, we adopt the following indirect approach: for each trio, we run PedMix on the child’s haplotypes to infer grandparents’ ancestry; we also run PedMix on the parents’ haplotypes to infer grandparents’ ancestry; we then examine whether the two inference results are consistent. That is, we view the inferred grandparents’ admixture proportions from parents as proxy of the true grandparental proportions. While this is not a direct evaluation, we believe it at least offers some hints on how well grandparental inference is likely to perform on real data.

We use the YRI and CEU populations as the two ancestral populations. Allele frequencies of these two ancestral populations and genetic maps from the HapMap project are used. We preprocess the data using the frequency-based trimming (default threshold is 0.4). We apply the phasing error removal procedure as described in the Methods Section. We use the reduced phasing error rate (see the Supplemental Materials for details) when running PedMix. Figure 9 shows the average admixture proportion inference error for parents and grandparents over ten trios. We show results for individual trios and also the average error. The average inference error for parents is about 3.4%, while the difference between the two inference results for grandparents is about 13%. Accuracy for most trios is relatively small, especially for parental inference. But for grandparent inference, the difference between the two estimates is relatively large for some trios. Figure 9 shows the parental/grandparental inference error with different amount of data (in terms of the number of chromosomes used). Our results show that inference error decreases when the amount of data increases, although sometimes the decrease in inference error is not very large. Overall, our results show that PedMix performs reasonably well on the real data.

**Figure 9:**
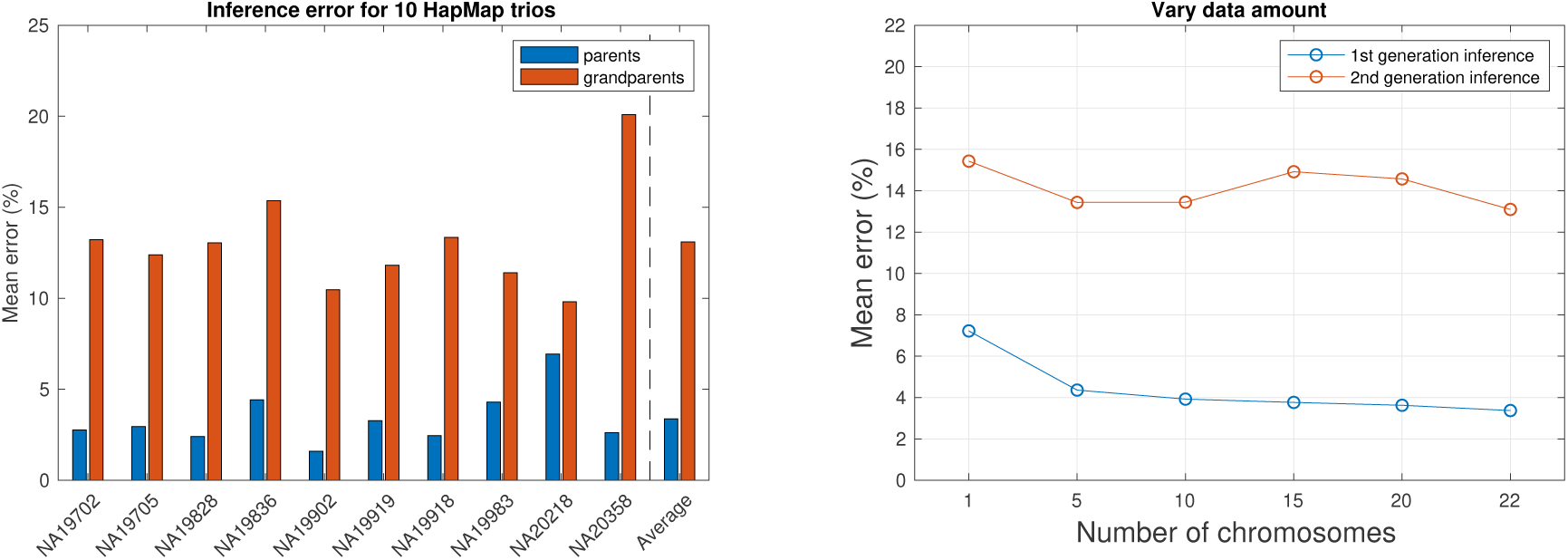
Admixture proportion inference error for parents and grandparents for ten HapMap trios from the ASW population. Left: inference error for ten HapMap trios individually or on average. Right: inference error with different amounts of data. X-axis: number of chromosomes used. Yaxis: parental/grandparental inference error. Parental: the difference between PedMix’s parental inference for the child and RFmix for the parents’ haplotypes. Grandparental: the difference between PedMix’s grandparental inference for the child and PedMix’s parental inference for the parents’ haplotypes in the trio.

## Discussion

In this paper we develop a method for inference of admixture proportions of recent ancestors such as parents and grandparents. To the best of our knowledge, there are no other methods for inferring admixture proportions for grandparents or great grandparents. The key idea is to use the distribution of admixture tracts which is influenced by the ancestral admixture proportions. We demonstrate that treating SNPs as independent sites is insufficient for inferring ancestral admixture proportions in general. See the Supplemental Materials for details. Our method uses a pedigree model, which is a reasonable model for recent genealogical history of a single individual. There are previous approaches for inferring ancestry that use additional information, for example geospatial information Yang, et al. (2014); Margalit,et al. (2015). Our approach only uses the genetic data from a single individual.

A natural question is whether our method can be extended to more distant ancestors. In theory it could, but as the number of generations increases, the amount of information for each ancestor decreases and the computational burden increases. This can be seen from Figure 5, where the inference error for great grandparents is significantly higher than those for parents or grandparents. On the other hand, Figure 5 shows that there are still information obtained from the inference even in the more difficult great grandparent case.

In Table 2, we show that PedMix can be used to infer the admixture proportions of an extant individual by averaging the inferred admixture proportions of ancestors. In comparisons with ADMIXTURE and RFmix, we find that the admixture proportions inferred from the average of ancestral admixture proportions in PedMix is comparable to that of RFmix and ADMIXTURE. The key difference between PedMix and RFmix/ADMIXTURE is that PedMix infers the admixture proportions of ancestors while the other methods only infer the admixture proportions of the focal individuals. Also, note that when we compare PedMix with RFmix and ADMIXTURE, PedMix uses recombination fractions in the founding populations, which are not used by RFmix and ADMIXTURE.

Inferences by PedMix are affected by the assumptions of the underlying population genetic processes (Figure 6). The inference error of PedMix can be significantly reduced if the recombination rate is high, admixture is more ancient, or the divergence time between the two source populations is large. Increasing the chromosome length has similar effect on inference accuracy as increasing the recombination rate.

In line with other studies (e.g. Anderson, et al. (2010)), we find that pruning of SNPs and preprocessing to remove potential phasing errors is critical for obtaining reasonably accurate results. In the Results Section, we compared two trimming strategies, LD pruning and frequency-based pruning. LD pruning is a common strategy used in HMM-based applications for removing background LD that is not modeled by the HMM. However, as low-frequency SNPs are more likely to have small values of *r*^2^, but are less informative for inference, strategies for removing SNPs based solely on measures of LD such as *r*^2^ might not be optimal. In fact, in the limited simulations performed here, we find perhaps surprisingly, that pruning strategies based on removing low-frequency SNPs, rather than SNPs in high LD, lead to the best performance. Based on our experience, we use the frequency-based trimming as our default data trimming approach. The objective of this paper is not to explore SNP pruning strategies for HMMs, but our results suggest that existing methods could be improved by devising better methods for SNP pruning.

PedMix works with haplotypes. At present, haplotypes are mainly inferred from genotype data and thus usually contain errors. Figure 7 shows that if untreated, phasing error can indeed greatly increase the inference error of PedMix. On the other hand, when we apply preprocessing to remove the obvious phasing errors, inference errors can be significantly reduced. Nonetheless, phasing error can still reduce inference accuracy. We note that phasing methods are constantly improving and the problem of phasing errors may be greatly reduced in the near future.

Simulation shows that PedMix can scale to whole genome data, when proper data preprocessing is performed. The current implementation of PedMix assumes two ancestral populations. In principle, PedMix can be extended to allow more than two ancestral populations, although this may lead to increased computational time.

## Supporting information

### S1 Supplemental Materials

Due to the space limit, we place parts of our results in the Supplemental Materials. These include additional methods, additional results and also a theoretical result which show that the ancestry inference problem studied in this paper will be infeasible without using linkage disequilibrium (LD).

## Acknowledgments

This work is partly supported by U.S. National Science Foundation grants IIS-1526415, CCF1718093 and IIS-1909425.

