## Supplemental Materials for "Inferring the ancestry of parents and grandparents from genetic data"

#### S1 Two strategies of data trimming

##### S1.1 Frequency-based pruning

The inference accuracy of our method largely depends on the amount of data. Recall that the computation of $h(AC_i)$ takes $O(N_K log(N_K))$ time for one site. Thus, large number of SNPs result in long computational time. We note that SNPs that have similar allele frequencies in the two source populations are less informative. Here, we use two simple thresholds to filter out less-informative SNPs to enhance the performance of our method.

1. Delete the SNPs that have zero or close to zero recombination fractions with their immediate neighboring SNPs.
2. Choose an allele frequency difference threshold $d_f$. Delete SNPs with population allele frequency difference between the two ancestral populations less than $d_f$.

##### S1.2 LD pruning

Before LD-pruning, rare variants with combined minor allele frequencies in the two ancestral populations lower than $f$ are removed. We use the correlation coefficient of linkage disequilibrium, $r^2$, in the ancestral populations to measure the level of linkage disequilibrium between two SNP sites. We scan through the SNPs sequentially. If $r^2 > c$ (with default value of $c$ is 0.1) between the current SNP and the previous SNP within a window of length $W = 10Kbp$, in either of the two ancestral populations, then the current SNP is removed.

#### S2 Preprocessing for phasing error

Phasing error results in a switching between two haplotypes, which is similar to how recombination affects haplotypes. The difference is that it only occurs in the current generation. Phasing error adds more noise in our model, especially when phasing error occurs much more frequently than recombination. Empirically, it is known the recombination rate for human is approximately $10^{-8}$ per generation between two adjacent base pairs. In most current data (e.g., haplotypes from the 1000 Genomes Project), phasing error occurs as frequently as once every 50 kb, which is three orders of magnitude larger than the recombination rate. So it is necessary to reduce the effect of phasing error. For this, we preprocess the haplotypes to reduce phasing error. Here are the steps that we use to remove likely phasing error for two extant haplotypes.

1. With the allele frequencies of two ancestral populations at each site, we first make a rough estimate of ancestry for the genotype $G$. For example, suppose the allele

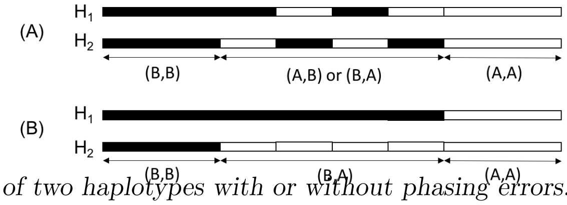


**Fig S1.** *Ancestry of two haplotypes with or without phasing errors. (A) Ancestry with phasing error. (B) Ancestry without phasing error. $H_i$ denotes the ancestry for haplotype $H_i$. Ancestry consists of blocks in white and black, where white denotes ancestral population A and black denotes ancestral population B.*

frequencies (of allele 1) for the two ancestral populations $A$ and $B$ are 0.1 and 0.8 respectively. It is more likely that two alleles (0, 1) at a SNP site have ancestry as $(A, B)$, and (1, 1) have ancestry $(B, B)$.

2. For each site of genotype $G$, we assign a "dominating ancestry". A dominating ancestry is an ancestry with one of these four possible pairs, $(A, A)$, $(A, B)$, $(B, A)$, or $(B, B)$, that appears most frequently within a region of certain length. Here we use the estimated number of SNPs between two phasing errors as the region length. We view $(A, B)$ and $(B, A)$ as type 1 ancestry, $(A, A)$ as type 0, and $(B, B)$ as type 2.

3. In the region that has the dominated ancestral type 1, we switch the two haplotypes if its ancestral painting $(A, B)$ or $(B, A)$ is different from its previous positions.

With the assignment of dominating ancestry, the two haplotypes phased from the genotype $G$ can be viewed as blocks of different dominating ancestry. For example, Figure **??** shows the ancestry for two haplotypes with black blocks indicating ancestry $A$ and white blocks indicating ancestry $B$. Figure **??** (A) provides an example on the ancestry of two phased haplotypes. Here we divide the whole region into three types of sub-regions: type 2 for $(B, B)$, type 1 for $(A, B)$ or $(B, A)$ and type 0 for $(A, A)$. Note that in type 0 and 2 regions, it is not obvious (and also not necessary) to detect and fix phasing errors. In a type 1 region, when we detect switch-overs between ancestry $(A, B)$ and $(B, A)$ within the region, we consider such switch-overs as the phasing error position and switch the suffix of two haplotypes from this point to make it consistent. This is because the probability that two recombination events happen at exactly the same place is $10^{-8} \times 10^{-8} = 10^{-16}$, which is much smaller than the phasing error probability of $10^{-5}$.

We note that the three-steps strategy described above does not remove all phasing errors. In fact, it may even add switching errors in some rare cases. However, our simulations show that this procedure can reduce a significant amount of obvious phasing errors, and help to reduce the noise of data (see the Results section). Without preprocessing, phasing error rate for genotypes is approximately 1 over 50kb, which is $p_p = 0.00002$. Preprocessing for phasing errors reduces approximately 2/3 phasing errors for admixed individuals. One can use a smaller phasing error rate $p_p = 0.0000066$ in PedMix after preprocessing.

### S3 Expected accuracy by random guess

In order to provide a baseline for the evaluation of the inference accuracy, we use a Bayesian model based random guess for estimating ancestry admixture proportions. Here we assume that the mean of admixture proportions of the ancestors is the admixture proportion of the focal individual. We treat each SNP position independently in the following. Given a genotype of a focal individual, we first sample ancestry for each SNP site based on the allele frequency of the SNP in the two ancestral populations. Ancestry of an allele (of some individual) refers to which of the two ancestral populations this allele originates from. With the sampled ancestry of this focal individual, we sample ancestry for his/her ancestors (parents, grandparents or great grandparents) following the posterior distribution. For example, the posterior probability of the parental ancestry is given in Equation **??**.

$$p(A_1, A_2|A_0) = \frac{p(A_0|A_1, A_2)p(A_1)p(A_2)}{p(A_0)} \quad (1)$$

Here $A_0$ is the sampled ancestry for one haplotype of the focal individual at a SNP site. $A_1$ and $A_2$ are the sampled ancestry of the two haplotypes from a single parent (which provides the allele for the focal individual). With the Mendelian segregation laws, $p(A_1) = p(A_2) = \frac{1}{2}$ are prior probabilities. The grandparental posterior probability $p(A_1, A_2, A_3, A_4|A_0)$ and great grandparental posterior probability $p(A_1, A_2, A_3, A_4, A_5, A_6, A_7, A_8|A_0)$ can be derived similarly. The estimate by the random guess is then computed using the sampled ancestry of each ancestor.

Note that random guess doesn't use information from admixture tracts and their lengths. For example, given the focal individual's genotype of pedigree CCCY with no phasing errors (Results Section ), The sampled ancestry of parents and grandparents all present ∼50% admixture proportions. Given genotype with no phasing error, random guess can still collect information for parents but fails to collect useful information for ancestors in grandparents and great grandparents. When adding phasing errors to genotype, random guess performs even worse in parents. For example, given the focal individual's genotype of pedigree CCYY (Results Section), random guess gets ∼50% ancestry for each parents while two parents are actually 100% and 0%.

### S4 Inference of ancestral admixture proportions with composite likelihood

The inference framework in PedMix takes advantage of the distribution of admixture tracts. Here, an admixture tract refers to a segment of the genome where the ancestral origin remains the same (i.e. coming from the same ancestral population). However, parental admixture proportions can be estimated simply from the distribution of genotype frequencies using composite likelihood as follows. Let $M = (m_1, m_2)$ be the admixture proportions of the two ancestors, then the composite likelihood is defined as the product of likelihoods in individual sites:

$$p(G|M) = \prod_i p(G_i|M) \quad (2)$$

The sampling probability for each site, $p(G_i|M)$, is calculated as a product of allele frequencies in the two parents using standard methods as follows. The probability of sampling an allele of type $j$ from a parent with admixture proportion $M$ and $1 - M$ from the population A and the population B respectively is $Mf_A + (1 - M)f_B$ if the allele frequencies of the allele $j$ at the site $i$ in the two populations are $f_A$ and $f_B$ respectively. We may infer admixture proportions by maximizing the composite likelihood. This can be done, for example, by performing a grid search over $M$.

This composite likelihood based method is computationally much faster than PedMix because it ignores linkage disequilibrium (LD). However, the method does not generalize to grandparents or more ancient ancestors as such models are not identifiable in the composite likelihood setting. To see this, let $M = (m_1, m_2, m_3, m_4)$ be the admixture proportions of the four grandparents, with $(m_1,\ m_2)$ being from one grandparental couple, and $(m_3,\ m_4)$ from the other. Let the allele $j$ be one of the two alleles of the genotype $G_i$. Without loss of generality, we further suppose this allele is from the parent (parent 1) descending from the grandparents with admixture proportions $m_1$ and $m_2$. The sampling probability, $p(G_i|M)$, is then obtained as a sum of products of terms like $p(j, allele\ from\ parent\ 1|m_1, m_2)$ by summing over both possible assignments of alleles to parents. Now,

$$
\begin{aligned}
p(j, allele\ from\ parent\ 1|m_1, m_2) &= \frac{1}{2}[\frac{1}{2}(m_1 f_A + (1 - m_1)f_B) + \frac{1}{2}(m_2 f_A + (1 - m_2)f_B)] \\
&= \frac{1}{4}((m_1 + m_2)f_A + (2 - (m_1 + m_2))f_B)
\end{aligned}
\tag{3}
$$

Equation **??** shows that $m_1$ and $m_2$ in $p(j, allele\ from\ parent\ 1|m_1, m_2)$ appear only as the sum $m_1 + m_2$. For any genotype $G_i$, the composite likelihood $p(G|M)$ only contains information about $m_1 + m_2$ but not $m_1$ and $m_2$ individually. Therefore, $m_1$ and $m_2$ are not separately identifiable in the composite likelihood model.

### S5 User guideline for using PedMix

Here is a list of user inputs needed by PedMix.

1. Phased haplotypes for the extant individual for whom we are to infer the admixture proportions of his or her ancestors. Haplotypes can be given in segments (or chromosomes), where segments are assumed to be independent.

2. For each SNP, allele frequencies of two ancestral populations.

3. Recombination fractions between adjacent SNPs along the haplotypes.

Often the input haplotypes may contain too many SNPs or less informative SNPs. In this case, the user may need to apply various data trimming techniques. We suggest to use frequency-based pruning with $d_f = 0.5$.

### S6 Additional results

#### S6.1 Details in comparison to ADMIXTURE, RFmix and ANCESTOR

##### S6.1.1 Verification

To verify that the average admixture proportions of ancestors of an individual provides a good estimate of the admixture proportion of focal individual, we simulate a sample of 100 individuals and compare the average values of admixture proportions of their ancestors with the admixture proportions of themselves. The sampled individuals are drawn either $g = 10$ generations or $g = 5$ generations after the time of admixture. The absolute difference (the error) between the true admixture proportions of one individual of the current generation and the average admixture proportions of his/her ancestors in the $K^{th}$ generation is computed as $|m^0 - \frac{1}{2^K}\sum_{1\leq j\leq 2^K} m^j|$, and we report the mean and variance of this as the mean error and the variance in the error among individuals (Table **??**). Here $m^0$ is the true admixture proportion of the sampled individual. $m^j$ is the true admixture proportion of the sampled individual ancestor $j$ in the $K^{th}$ generation.

**Table S1.** *The mean and variance (in unit of %) of difference between the admixture proportions of sampled individuals and the average of parental or grandparental admixture proportions.*

| Error (in %) | | $g = 10$ | $g = 5$ |
|---|---|---|---|
| Parental inference | mean | 0.567 | 0.151 |
| | variance | 3.98 | 5.45 |
| Grandparental inference | mean | 0.151 | 0.271 |
| | variance | 5.946 | 10.08 |

**Table S2.** *Pearson correlation coefficient for admixture proportion estimates from ADMIXTURE, RFmix and average of parents from PedMix.*

| correlation coefficient | ADMIXTURE | RFmix | PedMix (ave. of parents) |
|---|---|---|---|
| ADMIXTURE | 1 | 0.9975 | 0.9954 |
| RFmix | 0.9975 | 1 | 0.9945 |
| PedMix (ave. of parents) | 0.9954 | 0.9945 | 1 |

We find (Table **??**) that the average ancestral admixture proportions indeed approximately match the admixture proportions of the individual, as the mean differences are fairly close to 0 in Table **??**. Several aspects on the results in Table **??** worth attention. First, the variance in the error increases with more ancestors. Second, the variance tends to be larger for individuals from generations that are closer to the admixture event. This is because when the time since admixture is short, the individuals tend to have more different admixture proportions than individuals from a generation that is more distant from the admixture event and is thus well mixed.

##### S6.1.2 ADMIXTURE, RFmix and ANCESTOR setting

We apply ADMIXTURE and RFmix to infer the current generation admixture proportions on the same datasets. As suggested by ADMIXTURE, genotypes are preprocessed with LD pruning (see Section **??**) with parameters $c = 0.1$ and $W = 10Kbp$. To achieve the best performance in ADMIXTURE and RFmix, we also include all ancestral genotypes from the two ancestral populations along with 20 individuals from the admixed population to these two tools. That is, ADMIXTURE is run on "supervised mode". The number of ancestral populations $K$ is set to 2. ".bed" file is generated by PLINK. For real data, we use 170 haplotypes from CEU and 176 haplotypes from YRI as two ancestral populations in ADMIXTURE and RFmix to estimate admixture proportions for 61 genotypes from ASW. Here we compute the Pearson Correlation coefficient for admixture proportions estimates from ADMIXTURE, RFmix, and the average over parents by PedMix over 61 individuals in ASW population. The estimates by ADMIXTURE and RFmix show the highest correlation (0.9975, see Table **??**). The estimates by PedMix (average over parents) also have a high correlation with ADMIXTURE (0.9954) and RFmix (0.9945).

ANCESTOR infers the admixture proportions of parents of a focal individual given the ancestry state of each position in the genome. ANCESTOR allows phasing error in genotypes and can be used for multiple ancestries. In this paper, we use the ancestry inferred by RFmix from the Viterbi decoding as the ancestry states in ANCESTOR.

#### S6.2 Data trimming

To investigate the effect of frequency-based pruning and LD pruning, we perform a small investigation of the relative effect of LD-pruning and frequency-based pruning on the same simulated dataset. To efficiently compare the two trimming strategies, we simulate a shorter length genome $L = 5 \times 10^8$. The simulation settings are chosen in order to better compare the two ways of trimming and also to ease the computational burden.

There are $\sim 2.44M$ SNPs simulated for the whole genome. Here we examine two cases of data trimming. In both cases, we use a window size of $W = 10Kbp$ and $c = 0.1$ for LD pruning following the procedure described in Section **??**. In the first case of LD-pruning, we use $f = 0.05$ to remove rare variants (i.e., SNPs with combined frequency in the two populations being smaller than $f$), this results in 284K SNPs left. We compare frequency-based pruning with the threshold $d_f = 0.27$ to match the similar amount of SNPs (283K) in this case. In the other case, we remove rare variants with $f = 0.2$, and compare frequency-based pruning with $d_f = 0.5$, resulting in 97K and 94K SNPs respectively (Table **??**). In both cases inferences improve as more SNPs are removed. Overall the frequency-based pruning approach is slightly better than the LD pruning approach, at least in this simulation. We have, therefore, used frequency based trimming as the default strategy for data trimming.

To investigate the effect of frequency-based pruning, we simulate haplotypes for a small region of length $10^6 bp$ with 545,302 SNPs. The parameters used in the simulation are described in the caption of Figure **??** and we investigate different values of the previously explained allele frequency thresholds, $d_f$. Notice that trimming results in a substantial reduction in mean error, particularly for inferences of admixture proportions in grandparents. However, when the trimming threshold is too large, an increase in the mean error is observed due to the reduction in number of SNPs. In this case, the optimal trimming threshold appears to be around $d_f = 0.5$.

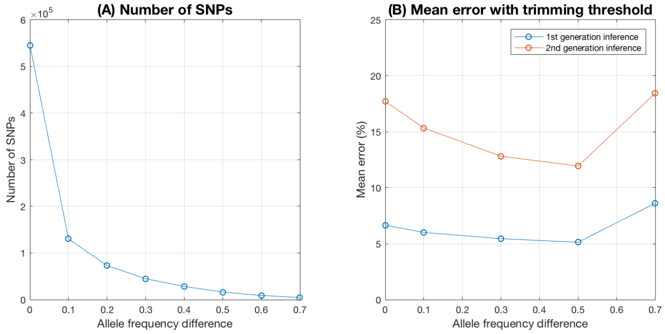


**Fig S2.** *Impact of data trimming threshold, $d_f$, on amount of data and inference accuracy. (A) The number of SNPs remaining after trimming. (B) The mean error of parent and grandparent admixture inference. Simulation parameters: $\mu = 10^{-6}$, $\rho = 5 \times 10^{-6}$, $nsam = 2,000$, $L = 10^6$*

As shown in Figure **??**, trimming using larger values of $d_f$ value results in a substantial reduction in mean inference error, especially for inferences of admixture proportions in grandparents. However, when the trimming threshold is too large (e.g. $d_f = 0.7$), an increase in the mean error is observed as too much information is now being lost by removing SNPs. The optimal trimming threshold depends on many parameters, such as mutation rate and the split time of the ancestral populations. It is challenging to determine the optimal value of $d_f$ for specific dataset. As a rule of thumb, it is desirable to have at least 100 SNPs per tract (from one ancestral population) after frequency-based pruning.

**Table S3.** *Compare the effects of LD pruning and frequency-based pruning on inference accuracy. LD: results with LD pruning. F-prune: results with frequency-based pruning.*

| Inference error (%) | LD (284K) | F-prune (283K) | LD(97K) | F-prune (94K) |
|---|---|---|---|---|
| parents | 14.99 | 14.38 | 13.03 | 7.93 |
| grandparents | 18.02 | 16.48 | 16.05 | 14.79 |

Clearly, more work is needed to identify optimal pruning strategies for real data analyses, for the current method and for other methods that use HMMs for population genetic inferences. However, such investigations are not the main subject of the current paper.

##### Impact of data amount

We run PedMix on different amounts of data, by sub-sampling 5, 10, 15 and 20 chromosomes, to evaluate the effect of data amount on inference accuracy. The mean error is estimated over 10 samples. From Fig **??**, we can see a clear linear decrease of mean error for parents, grandparents and great grandparents inference, as more data are added. The highest mean error for five chromosomes is 18.07%, which is still much lower than the random guess (about 35%).

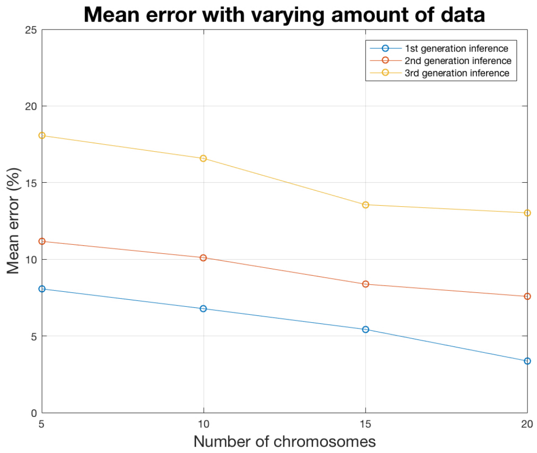


**Fig S3. Impact of data amount** We use 5, 10, 15, and 20 chromosomes respectively. For great grandparents inference, $d_f = 0.9$ is used due to computational burden.

#### S6.3 Impact of recombination rate on a larger genome

As discussed in the main text, the mean error for parental and grandparental inferences decreases as recombination rate increases. Results are shown in Figure **??**. As we simulate genomes with longer length, the increase of recombination rate does not lead to the significant decrease in mean error. That is, the mean error asymptotes as recombination rate increases. This happens especially when the simulation length is long (at the default setting $L = 3 \times 10^9$), the decrease of mean error is only about 1% as recombination rate increases 10 times.

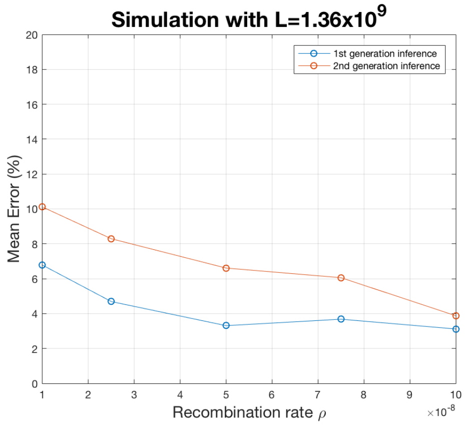


**Fig S4.** *Varying recombination rate using 10 chromosomes*

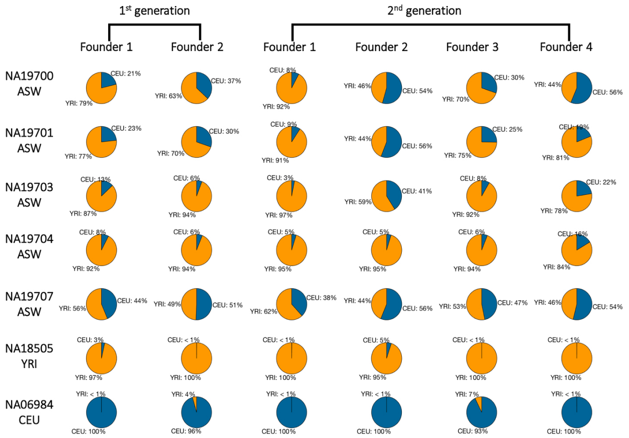


**Fig S5.** *Example pie charts of admixture proportions for 6 individuals. For each individual, we display four pie charts which represent four grandparents respectively. The blue part shows the percentage of CEU population and the orange part shows the percentage of YRI population.*

#### S6.4 Results on the 1000 Genomes Project data

We run PedMix on the data from the 1000 Genomes Project . The 1000 Genomes Project recently released phased haplotypes on 22 chromosomes for 1,092 individuals. We analyze data from the CEU (Utah Residents with Northern and Western European Ancestry), YRI (Yoruba in Ibadan, Nigeria) and ASW (Americans of African Ancestry in SW USA) populations. The African-American population tends to have admixed European and African (primarily West-African) ancestry. Here we regard CEU and YRI as the two source populations for ASW and infer the admixture proportions of parents and grandparents of ASW individuals. For these three populations, there are 85 CEU individuals (170 haplotypes), 88 YRI individuals (176 haplotypes) and 61 ASW individuals (122 haplotypes) in total in the 1000 Genomes Project data.

We approximate the allele frequencies in the two hypothetical source populations using the average allele frequencies in the CEU and YRI populations. The original data has 1,060,387 SNPs in total. After applying the frequency-based pruning with $d_f = 0.5$, there are 256,122 SNPs (about 24%) left. The recombination fractions are calculated

**Table S4.** *Comparison of admixture proportion estimate among ADMIXTURE, RFmix and PedMix for five ASW individuals. The PedMix results are the averages of the admixture proportions of parents and grandparents inferred by PedMix. These averages are used as proxies for focal individuals, and are then compared with estimates of ADMIXTURE and RFmix. The percentage of CEU origin is shown in table.*

| individual ID | ADMIXTURE | RFmix | PedMix (parents) | PedMix (grandparents) |
|---|---|---|---|---|
| NA20296 | 22.06% | 21.78% | 20.46% | 17.37% |
| NA19625 | 32.02% | 34.35% | 31.96% | 31.13% |
| NA19921 | 41.22% | 41.20% | 42.61% | 39.84% |
| NA20414 | 63.84% | 68.76% | 63.88% | 67.00% |
| NA20314 | 92.38% | 97.93% | 88.76% | 88.13% |

based on the recombination hotspot map of the 1000 Genomes Project. We sample 7 individuals from the CEU, YRI and ASW populations and apply PedMix to infer admixture proportions in their parents and grandparents.

The inference results of admixture proportions for parents and grandparents for seven 1000 Genomes individuals are shown in Figure **??**. We infer the admixture proportions of the CEU individual ancestors to be 98% of CEU origin on average. Similarly, the admixture proportions of YRI individuals ancestors are 98% of YRI origin on average. The admixture proportions in the African-American ASW population vary considerably among individuals (Figure **??**). Note that some ancestors for the CEU and YRI individuals have small (but non-zero) inferred admixture proportions. Since the proportions are very small (within the error margin of our method), we cannot determine whether these ancestors are admixed or not.

To further validate our results, we analyze 61 individuals from ASW population using ADMIXTURE and RFmix. Genotypes are pruned with the default LD pruning setting (see Section **??** of the Supplementary Material). Meanwhile, genotypes from CEU and YRI populations are provided as the two ancestral populations in both tools. Using PedMix, we compute the average admixture proportions from parents and grandparents to see if the results are consistent with those from ADMIXTURE and RFmix. The percentages of the CEU origin of the five ASW individuals are shown in Table **??**. We list five individuals from ASW with different level of admixture proportions (from 20% to 90%). Although the true admixture proportions of these individuals are unknown, the results of PedMix are consistent with those from RFmix and ADMIXTURE. Moreover, the estimate by PedMix is highly correlated with those by ADMIXTURE and RFmix. Pearson correlation coefficients are 0.9954 and 0.9945 respectively (Table **??**).

#### S6.5 Running Time

We now evaluate the computational efficiency of PedMix. PedMix is written in C++. To make the algorithm run faster, we not only adopt the divide-and-conquer strategy, but also make it run with multi-threads. Multi-threading can be useful when there are multiple chromosomes in the data. The best performance occurs when there are $k$ chromosomes with similar number of SNPs using $k$ threads in parallel. However, since it is an optimization problem, the convergence time is uncertain. In general, the running time increases exponentially with the number of generations inferred. Here we report the average running time of grandparent inference for 10 individuals and fix the number of threads to 1 as we increase the number of SNPs from 5,000 to 550,000 (Figure **??**). As expected, we observe a clear increase in time when we use more SNPs.

For comparison, ADMIXTURE, RFmix and PedMix are run on same datasets in Results Section. To estimate admixture proportions over 20 individuals, ADMIXTURE

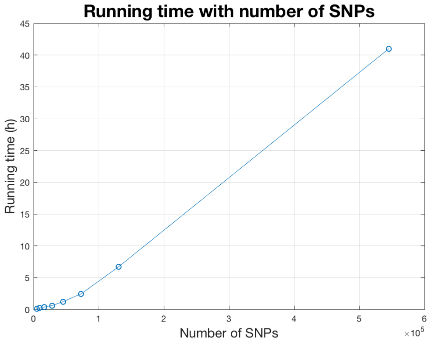


**Fig S6.** *Running time with various number of SNPs using one thread.*

takes 1.5 min using 20 threads, while RFmix takes 21.5 mins using one thread. On average, PedMix takes 3 mins for parental inference and 2.5 hrs for grandparental inference using 11 threads for each individual. ADMIXTURE and RFmix run much faster than PedMix. This is because the computation performed by ADMIXTURE and RFmix and the parameters to estimate are very different with those of PedMix.
